# *Hydra* in Flow – Investigating the Flow Response of *Hydra* via Microfluidic Systems ^*^

**DOI:** 10.1101/2025.08.30.673271

**Authors:** Ena Ysabella D. Arroyo, Jaimie A. De Guzman, Maro M. Aziz, Helen Cardenas-Rios, Roquel John Roman, Robert E. Steele, Siavash Ahrar

## Abstract

*Hydra*, a simple freshwater cnidarian, occurs in both moving and still bodies of water. Flow and corresponding forces are ubiquitous factors in *Hydra*’s environment. Even in still water, brief exposure to a burst of flow (e.g., due to wind) can influence the surface attachment and dispersal of polyps. Additionally, alignment with flow may play a crucial role in minimizing forces and regulating feeding behaviors in *Hydra*. However, the response to flow (particularly in the presence of gravity) has remained underexplored. Using vertically oriented microfluidic chambers, we investigated the biomechanical response of *Hydra vulgaris* and two additional *Hydra* species to fluid flow. For *Hydra vulgaris*, strong surface attachment was observed for all flow rates (0 - 100 mL/hr, corresponding to average velocities of 0 - 2.2 mm/s for the chambers). The experiments indicated alignment of the body column with the flow direction at high flow rates (≥ 50 mL/hr). Alignment with flow was examined by quantifying the angle between the organisms (using a vector connecting head-to-foot) and the flow direction for multiple individuals (N = 9). Most individuals of *H. vulgaris* exhibited alignment under high flow. While preliminary, comparisons with *H. hymanae* and *H. oxycnida* suggested species-specific differences *H. hymanae* showed tentacle deformation but lacked clear alignment. *H. oxycnida* did not remain attached under flow. Additional experiments explored the combined effects of flow and osmolarity. These results support the use of microfluidic tools to examine flow-related behaviors and highlight the potential for comparative biomechanics across *Hydra* species.

## 1 Introduction

The freshwater cnidarian *Hydra* occurs in both moving (rivers and streams) and still bodies of water (vernal pools, ponds, and lakes) on all continents except Antarctica (Martínez et al., 2010 [1]). Even in still water, short-term exposure to a burst of flow (e.g., due to wind) could influence the polyp’s survival. Specifically, exposure to flow could interrupt surface attachment or change body orientation, influencing dispersal or feeding behaviors. Investigations in sea anemones (e.g., *Metridium senile*) indicate that a change in shape leads to a dramatic change in the drag coefficient and the corresponding drag force (Koehl 1982[2]). In her investigation of *M. senile*, Koehl described the relationship between transport (maximizing food capture) and minimizing mechanical forces (Koehl 1977)[3]. Despite the ubiquity of flow in *Hydra*’s habitat, examining the interactions between flow, surface attachment, and body orientation has remained limited. This gap is in part due to the absence of tools that provide flow and imaging. In this effort, we sought to address this gap and investigate the flow response of *Hydra* in vertically-positioned chambers. Our choice was informed by an investigation that indicated a role for gravity (negative gravitotaxis), particularly for young buds (Ewer 1947)[4]. To this aim, we combined microfluidic chambers and custom microscopy systems to investigate *Hydra*’s flow response.

To interpret *Hydra*’s responses to flow, it is helpful to consider its basic anatomy and physiological mechanisms. *Hydra* has a simple body plan consisting of three regions: the foot, the body column, and the head (Vogg et al., 2019)[5]. The foot enables attachment to a surface using secreted adhesives (Rodrigues et al., 2016)[6]. The head region consists of a dome-shaped structure called the hypostome, surrounded by a ring of tentacles. *Hydra* lacks a permanent mouth. Instead, it transiently forms an opening at the apex of the hypostome for feeding (Campbell, 1987; Carter et al., 2016)[7, 8]. *Hydra* is composed of two epithelial layers, ectoderm and endoderm, separated by an extracellular matrix called mesoglea. In the absence of a centralized brain, behaviors are controlled by diffuse and regenerative nerve nets (Szymanski and Yuste, 2019; Keramidioti et al., 2024) [9, 10], one in each epithelial layer (Dupre et al., 2017; Keramidioti et al., 2024)[11, 10]. Mechanical behaviors are generated through epitheliomuscular activities and the regulation of hydrostatic pressure within the fluid-filled hydrostatic skeleton (Wang et al., 2023)[12]. Water enters the gastrovascular cavity via osmosis through the epithelial layers. To prevent overaccumulation of fluids, *Hydra* undergoes periodic body contractions, which expel the excess water through the mouth (Kücken et al., 2008; Yamamoto et al., 2020; Ferenc et al., 2021)[13, 14, 15]. Body contractions may also play a role in the maintenance of the microbiome (Nawroth et al., 2023)[16]. Behaviors are controlled by the nerve nets (Szymanski and Yuste, 2019; Keramidioti et al., 2024) [9, 10]. Sensory inputs are detected along the entire body. However, a recent study highlights that the head region may be critical in coordinating motor responses (Badhiwala et al., 2021)[17]. Despite its anatomical and neural simplicity, the organism exhibits complex and state-dependent behaviors (Yamamoto et al., 2023; Kim et al., 2024)[18, 19]. These behaviors are distinct and well-characterized (Han et al., 2018)[20]. We focused on how short-term (15-minute) exposure to flow and associated mechanical forces influences behaviors. To provide context, we review the existing microfluidic platforms for *Hydra* research.

Microfluidic systems are commonly used to study aquatic organisms. However, their application to *Hydra* research has remained limited. Badhiwala et al. developed resources, including an immobilization chamber for electrophysiology using embedded electrodes, a perfusion chamber for constrained locomotion, and a quasi-2D plane behavioral chamber (200 to 600 *µ*m in height) for long-term imaging (Badhiwala et al., 2018)[21] of *Hydra*. In a subsequent study, these authors developed a system that combined immobilization and pressurized microvalves (to provide well-regulated local stimuli) for functional recording of neuronal activity in response to touch (Badhiwala et al., 2021)[17]. Microfluidic channels have been used to study phototaxis and temperature regulation (Tzouanas et al., 2021; Kim et al., 2024)[22, 19]. Beyond whole animals, Perros and coauthors demonstrated an on-chip micropipette system to measure the mechanical properties of tissue spheres (Perros et al., 2024) [23]. We previously developed a light-sheet-compatible microfluidic chamber that enabled imaging of freely moving *Hydra* within millimeter-sized chambers (Hedde et al., 2023)[24]. During these preliminary investigations, we observed the alignment of the body column with the direction of an imposed flow. Although not performed using microfluidic systems, a key investigation by Khetan et al. estimated the forces due to flow that are needed to detach *H. vulgaris* and *H. magnipapillata* from surfaces with various material properties (Khetan et al., 2019)[25]. In this investigation, we used vertical microfluidic chambers (flow against the direction of gravity) for short-term (15 minutes) response of *Hydra* to flow.

We postulated that even in still waters, *Hydra* may experience brief exposures to strong flow (e.g., due to wind). Therefore, flow and corresponding forces could play an important role in the organism’s behaviors. In this report, we focused on surface attachment and alignment with the direction of flow.

## 2 Materials and methods

### 2.1 Microfluidic chambers

Chambers were developed as previously described (Hedde et al., 2023)[24]. Designs were laser cut (40 W laser cutter) from 1.6 mm acrylic sheets (matching the chamber depth). The majority of the experiments were conducted on straight channels that were 4 mm or 6 mm wide and 5 cm long. These plastic parts were attached to a flat substrate (plastic or glass slides) to construct the mold. Molds were then used for silicone casting (Polydimethylsiloxane or PDMS, Dow Sylgard™ 184 Silicone) using conventional protocols (1:10 curing agent to the base mass ratio). Molds were heated at 85°C on a hotplate until the PDMS was cured. The cured PDMS was cut from the mold with a craft knife. Inlets and outlets were created with a 2 mm diameter biopsy punch. These silicone parts were attached to glass slides using a plasma treatment (Harrick Basic Plasma Cleaner). The devices were further baked on a hotplate (95°C for two hours). Before the first use, the devices were placed inside a crystallizing dish containing water on a hot plate and boiled for an hour at 100°C to remove any potential contamination. After each experiment, devices were thoroughly rinsed with 70% ethanol solution, rinsed with water, and again boiled for at least an hour at 100°C to prevent potential cross-contamination (bioadhesive remaining from previous *Hydra*). Devices were typically discarded after the third experiment, primarily due to scratches on the glass slide or delamination of the silicone. Dimensional characterization of the system (i.e., Reynolds numbers) is provided in Supplementary Table 1.

### 2.2 Imaging system

We used our recently developed approach (Obenshain et al., 2024)[26] to visualize the vertically positioned chambers. Briefly, a USB camera was controlled using a Raspberry Pi 5 for image acquisition. The optical path included objectives (2x or 4x) and a tube lens (50 mm). For preliminary experiments involving fluorescence, emission filter (Chroma ET605/70m) was included in the optical path. However, the majority of the experiments were conducted with bright light with brightfield illumination from a Diamond Art Club Light Pad (Figure 1A). The chambers were secured on a 3D printed frame and secured in an upright (i.e., portrait) profile on an XYZ stage. The imaging components were mounted on a Z stage. These components were attached to an optical breadboard during the experiment.

**Figure 1:**
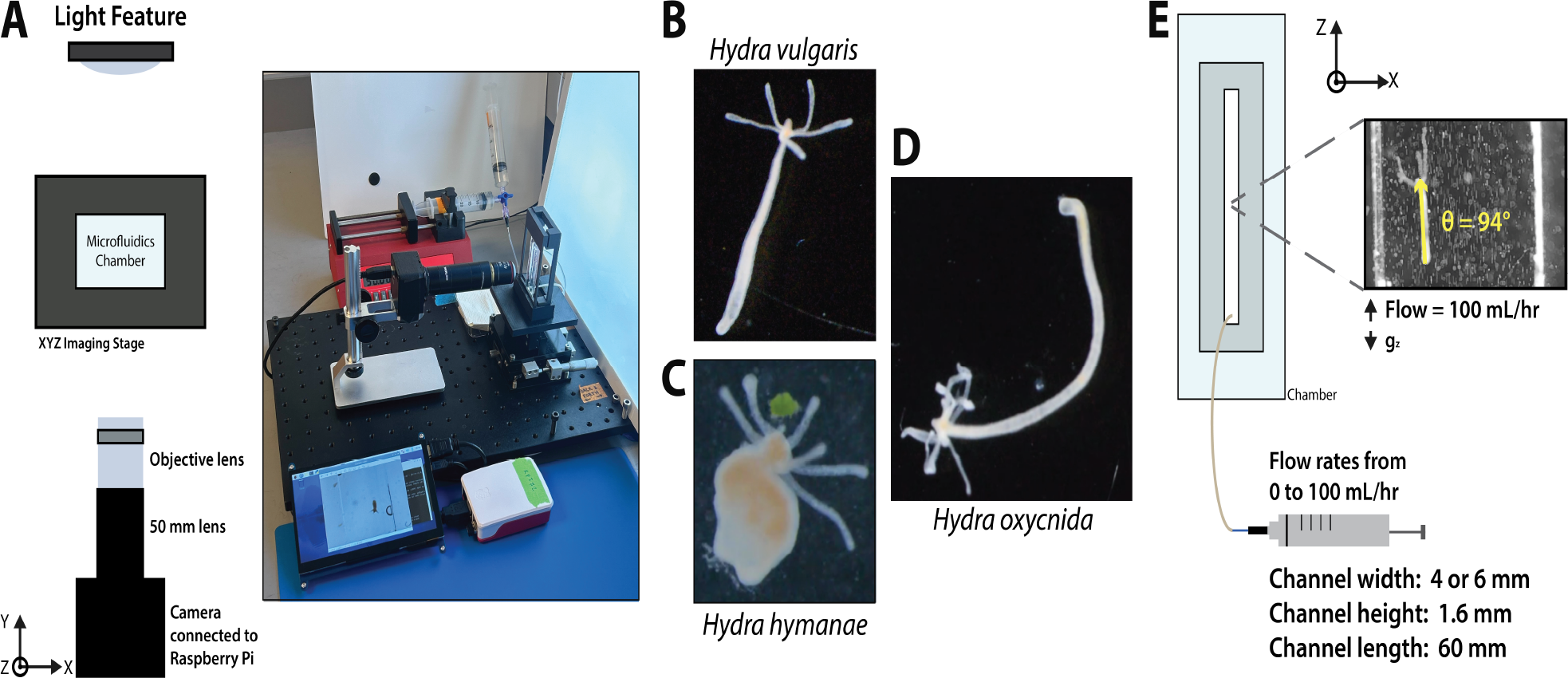
*Hydra* in flow. (A) Microfluidic chambers and do-it-yourself microscope were used to investigate the flow response of *Hydra* in vertically-positioned chambers. Polyps of (B) *Hydra vulgaris*, (C) *Hydra hymanae*, and (D) *Hydra oxycnida* were studied. *Hydra vulgaris* was the primary organism examined. (E) Chambers were positioned vertically in a portrait profile. Flow was introduced from the lower inlet against the force of gravity. To examine the alignment of the body column with the flow angle between the foot and head, measurements were made. Here, *θ* = 90°C represents perfect alignment with the direction of flow. The direction of gravity is highlighted with *g*_*z*_.

### 2.3 *Hydra* culture

*H. vulgaris* (Figure 1B) was the primary organism in this study, as it is the *Hydra* species most commonly used for experiments by other researchers. While anatomy is consistent in the genus, there is among-species variation in body size. For example some species have a stouter appearance since they remain partly contracted (*H. hymanae*, Figure 1C) [27] or have a larger body size (*H. oxycnida*, Figure 1D). Polyps were grown at room temperature inside crystallizing dishes containing *Hydra* medium. They were fed nauplii of *Artemia franciscana* once a week. For flow experiments, spring water (Aquafina™) was used. For osmolarity experiments, DI water for 0 Osmo, or a 50 mM sucrose solution (inside DI water) was used. The total durations of experiments with DI water were less than an hour since DI water negatively impacts *Hydra*.

### 2.4 Flow response

Using microfluidic chambers, polyps were exposed to volumetric flow rates (typically, 0 to 50 mL/hr) against the direction of gravity. Supplemental Figure 1 demonstrates a sample experiment with an adult polyp of *H. vulgaris* exposed to volumetric flow rates from 0 to 100 mL/hr. To quantify the alignment behaviors, we defined the vector between the polyp’s foot (i.e., the location of attachment to the glass) and its head. We used the vector to measure (a) the alignment of the body column with the direction of the flow, (b) the change in length of the body column (body length normalized to the initial value). The angle (*θ*) defined between the body vector and the horizontal cross-section of the chamber was used to quantify the alignment. Therefore, *θ* = 90° indicates perfect alignment with the direction of the flow. The angles were measured typically every 15 seconds, with the entire recording lasting 15 minutes for each condition. The order of flow rates were randomized at the start of each experiment. A rest period of at least 2 minutes was provided between the test conditions. We also examined the effects of osmolarity and channel shape on *H. vulgaris*. These preliminary results demonstrate the utility. All experiments were conducted on vertically-positioned chambers with and without flow.

### 2.5 Data analysis

Image analysis, particularly angle measurements, was performed using Fiji (Schindelin et al., 2012)[28]. Circular data analysis was performed using the True Circular Mean [29] and the Matlab Toolbox for Circular Statistics[30].

## 3 Results

The experiments with adult polyps of *H. vulgaris* led to the following observations. First, the polyps remained surface attached to the glass for all volumetric flow rates (corresponding to surface shear stresses ranging from 0 to 8.2 mPa - see supplementary Table 1). Without flow (0 mL/hr) or in low flow (1 mL/hr) the polyps’ head typically pointed towards the inlet, perhaps due to gravity. Increasing the flow rate to 10 mL/hr led to more exploration of the chamber by various movements including swaying and bending. At high flow (50 mL/hr) the body column was aligned with the direction of the flow (as measured by the angle), with the head pointing away from the inlet. Figure 2 demonstrates the alignment results as polar plots for three independent polyps, where the angle *θ* = 90° represents perfect alignment with the direction of the flow (vertical against gravity). The size of each vector is normalized to the initial length of the polyp. Next, the circular mean and circular standard deviation were calculated for each condition for all polyps. Figure 3 provides a summary of all experiments. In Figure 3A, each color represents the circular mean of a polyp for all flow conditions. The results indicate that the majority (6 out of 9) polyps were aligned with the flow at high flow rates. We also calculated the weighted circular mean and standard error of the mean for each flow rate (Figure 3B). Supplemental Table 2 summarizes the results both at the individual polyp and overall levels. Using V-tests we examined the alignment for each flow condition (using the 9 circular means for a given flow rate). Results indicated an alignment with the direction of flow (identified as *θ* = 90°) for the high flow rate (50 mL/hr) condition (p = 0.0004). V-test results indicated that other flow rates were not aligned with the direction of imposed flow.

**Figure 2:**
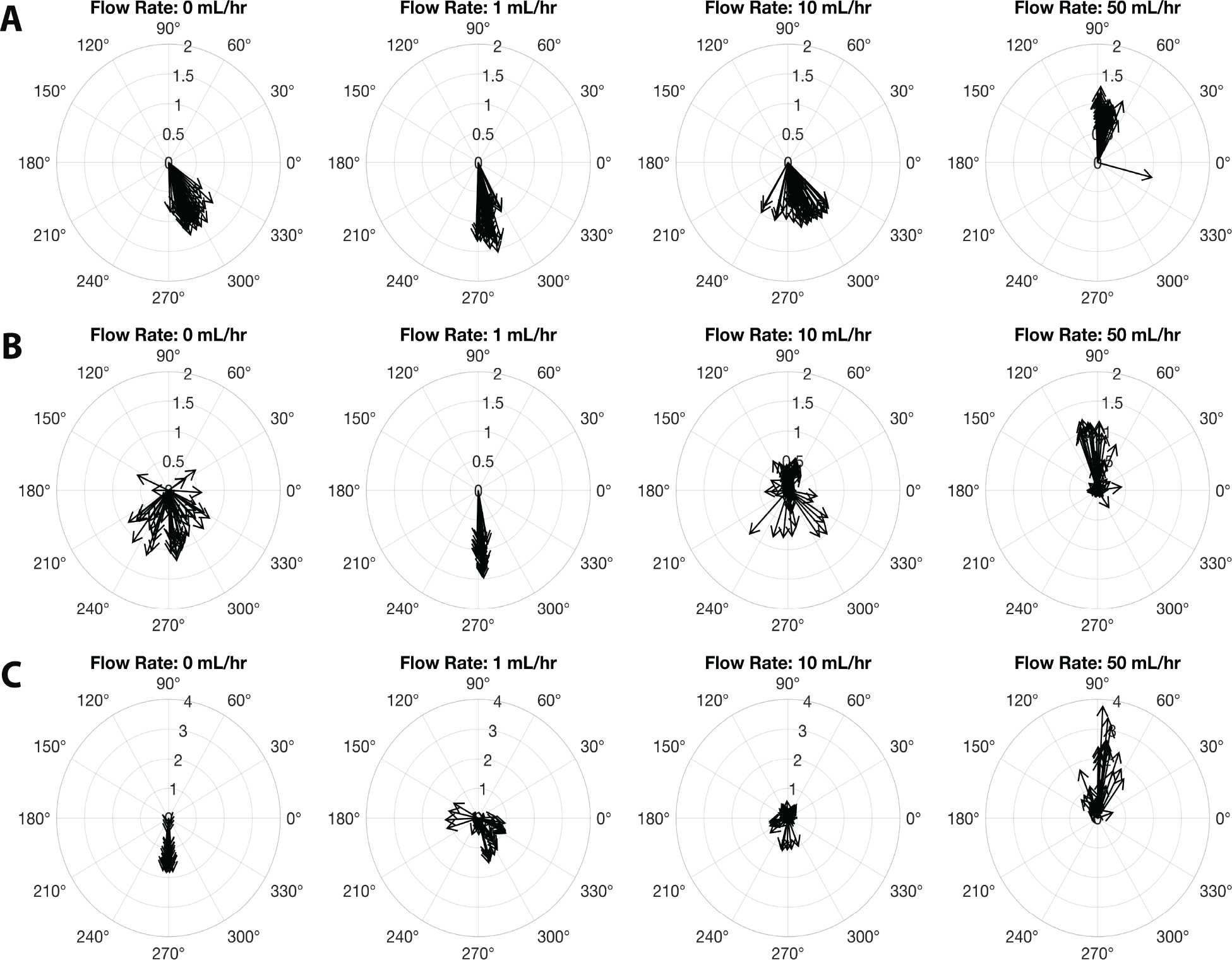
Three polyps in flow. Three sample polyps indicate the alignment with the direction of the flow at 50 mL/hr. Each row represents a single *Hydra vulgaris* polyp under the four flow conditions. The size of the vector represents the size of the polyp normalized to the initial size. For each condition, the polyps were exposed to flow for 15 minutes. Angles and polyp size were measured every 15 seconds for the entire recording. The order of flow conditions was randomized at the start of each trial.

**Figure 3:**
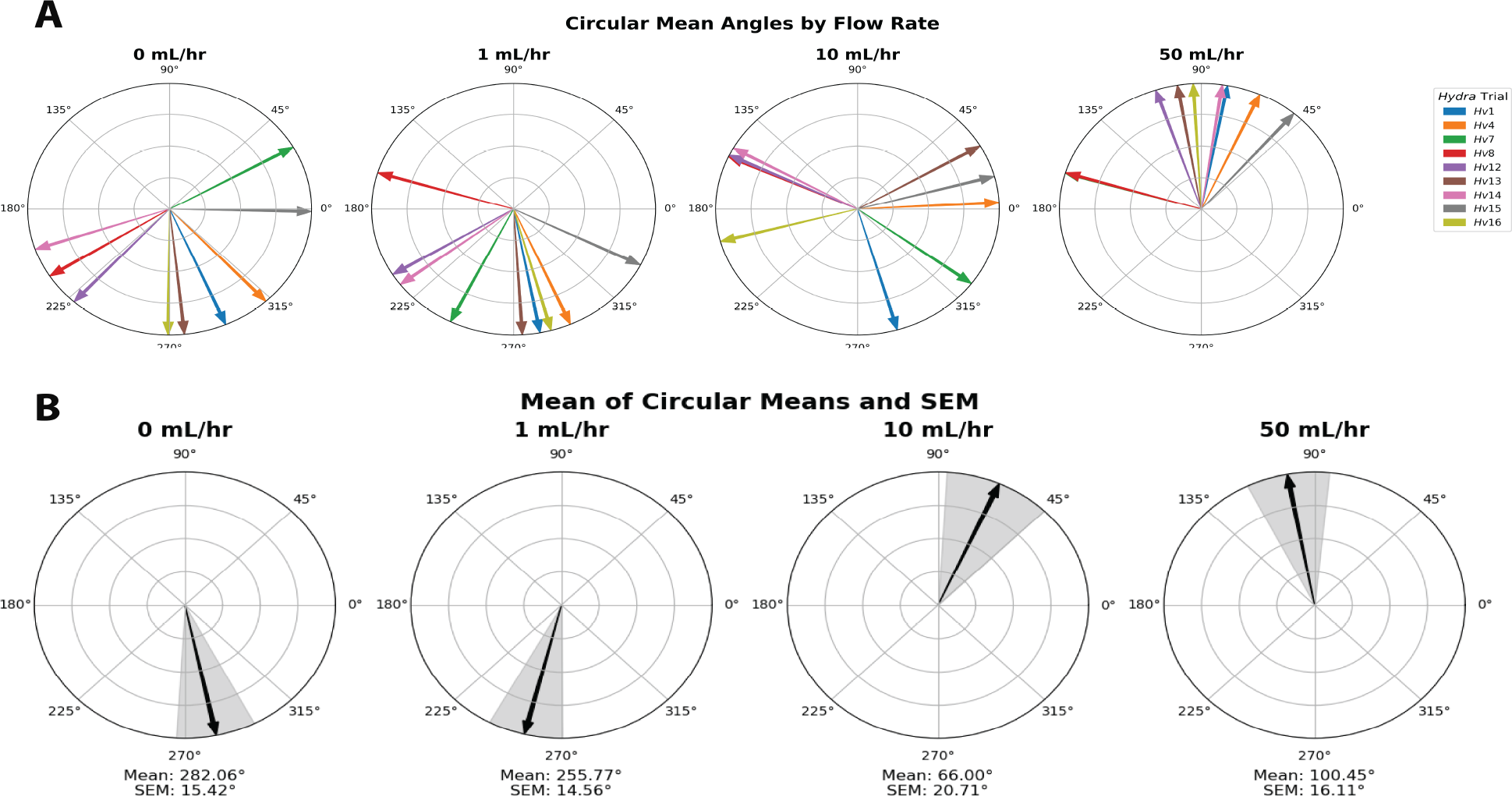
Assessment of alignments with flow as depicted by circular means for each polyp. (A) For each polyp the circular means for all flow conditions were calculated. At no flow or low volumetric flow rates (i.e., 1 mL/hr) most organisms point towards the bottom of the chamber. At high flow rates (i.e., 50 mL/hr), 6 out of 9 organisms are in the direction of the flow. (B) The aggregate circular means for 0 and 50 mL/hr and standard error of the mean (SEM). A V-test against *θ* = 90 ° indicated that the polyps were aligned with flow at 50 mL/hr. Table 2 provides a summary of results including circular standard deviations.

Next, we examined the polyps of *H. hymanae* (Figure 1C) and *H. oxycnida* (Figure 1D), under these same flow conditions. *H. hymanae* had similar surface attachment to *H. vulgaris* under vertical flow. However, given the contracted body structure, there was no clear alignment with the flow for the conditions examined. For *H. hymanae*, we observed retraction and bending of the tentacles (Figure 4) under flow. In contrast, *H. oxycnida* attached to surfaces in no flow, but did not remain surface attached in the presence of flow at any velocity in the chambers (Supplementary Figure 3). In the presence of flow, the organisms were washed away.

**Figure 4:**
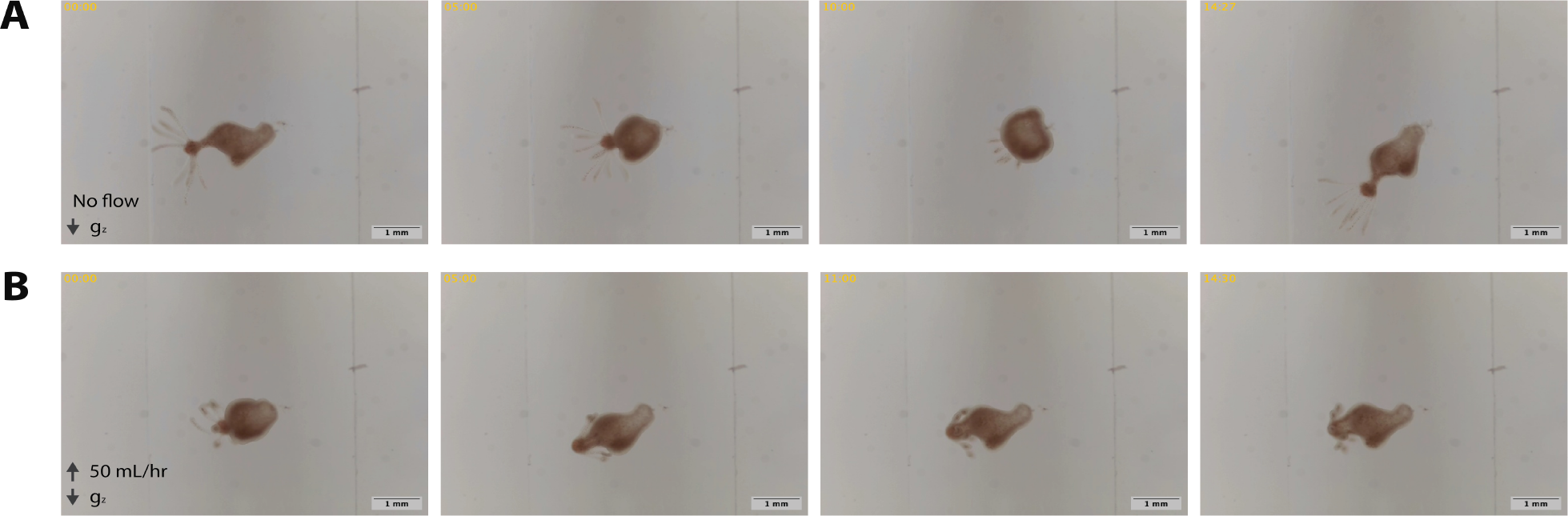
*Hydra hymanae* response to flow. *H. hymanae* has a contracted body column compared to *H. vulgaris* (A) Under no flow conditions the organism demonstrated swaying and contractions. The tentacles were elongated. (B) With flow conditions, the tentacles were bent. However, at the flow rates tested no alignment with the flow was observed. For these experiments the polyps remained surface attached.

**Figure 5:**
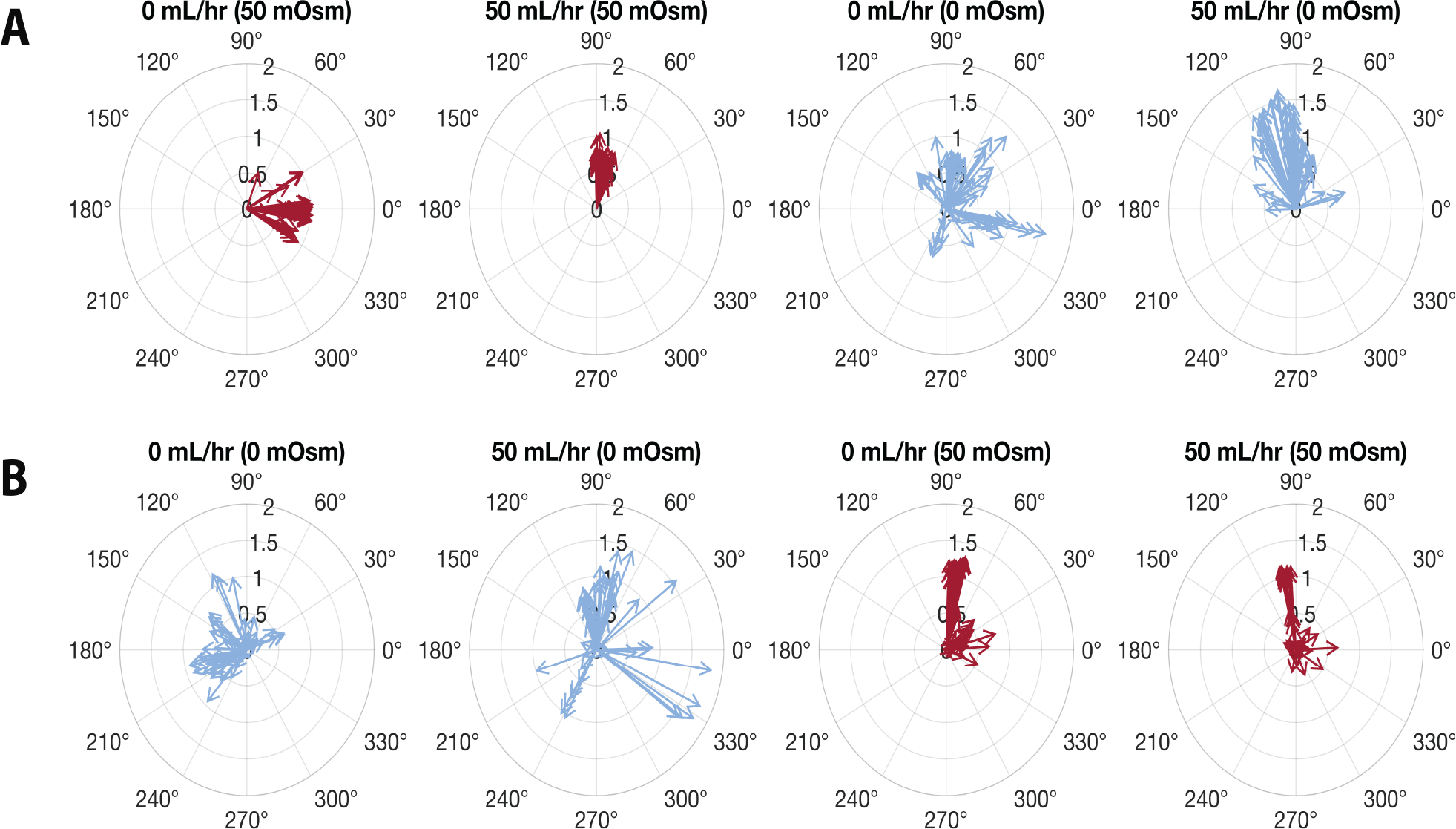
Flow and osmotic pressure. Two polyps were exposed sequentially to 0 and 50 mL/hr flow of 0 Osm/L (for DI water) or a 50 mOsm/L (50 mM sucrose) solutions. To ensure that there were no effects due to the order of the solutions, both high-to-low and low-to-high osmolarity sequences were examined. Strong surface attachment was observed for both conditions. (A) High to low osmolarity tests. With imposed flow for both osmolarity conditions there were alignment with the direction of the flow. At high osmolarity conditions, the overall movement was limited as compared to low osmolarity conditions. (B) Low to high osmolarity tests. There was some alignment with the direction of flow for both osmolarities.

**Figure 6:**
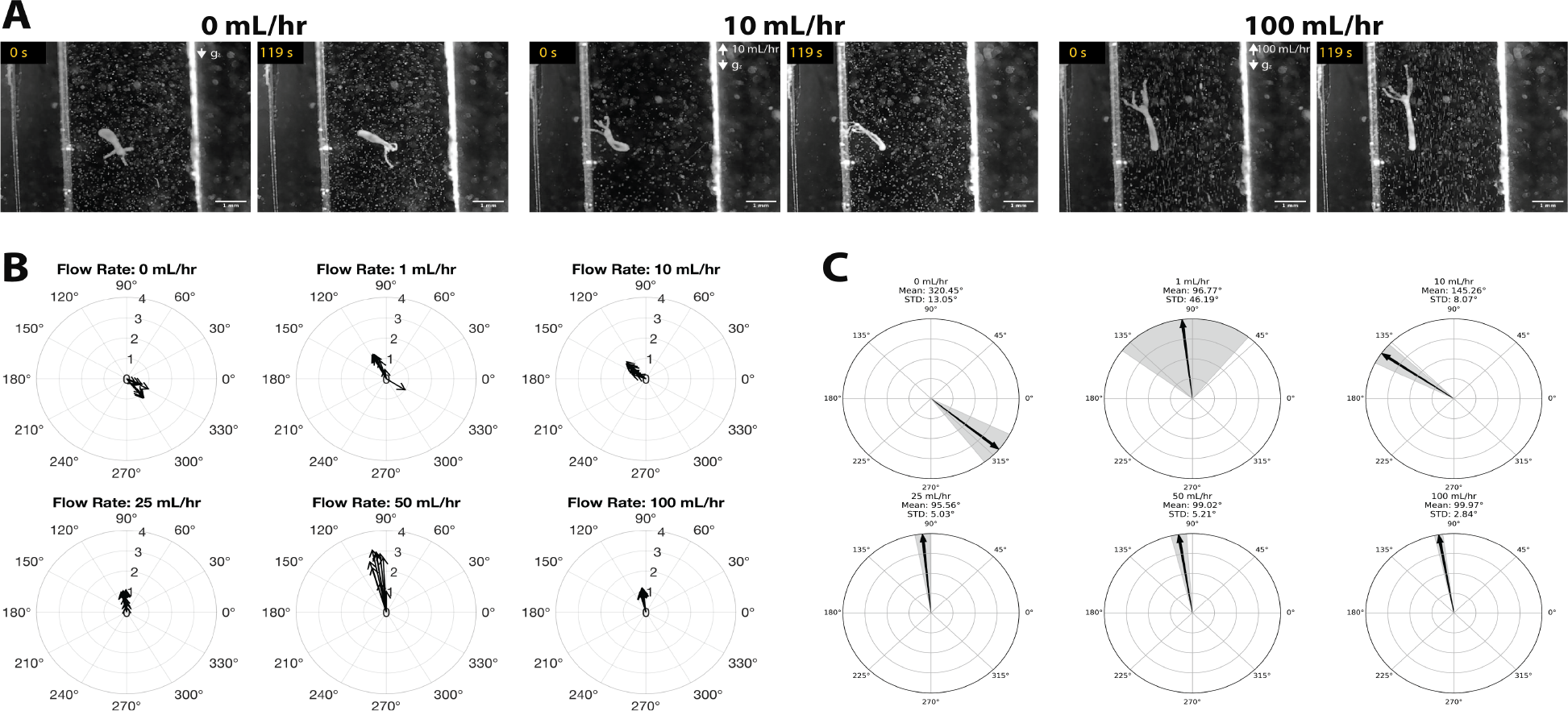
Sup Figure 1: *H. vulgaris* polyp in flow (0 - 100 mL/hr). A polyp was exposed to 0, 1, 10, 25, 50, and 100 mL/hr volumetric flow rate (each condition was for 2 minutes). In this experiment, microbeads were used for flow visualization. (A) Starting and final position of the polyp for 0, 10, 100 mL/hr. (B) Polyp’s normalized size and orientation relative to the flow (measured every 10 seconds from the entire recording). When exposed to low volumetric flow rates 0 - 10 mL/hr there was poor alignment with the flow throughout the duration. Specifically, at 0 mL/hr, the polyp was positioned towards the bottom of the channel, possibly due to gravity. At higher volumetric flow rates (25 mL/hr and above), the polyp aligned with the direction of the flow. The size of the vector represent the size of the polyp normalized to the initial size. (C) Alignment with flow were quantified from the measurements using the circular mean and circular standard deviation for each condition. There were noticeable alignments with flow starting at 25 mL/hr. The standard deviation of the angles decreased with the increase in flow rate.

**Figure 7:**
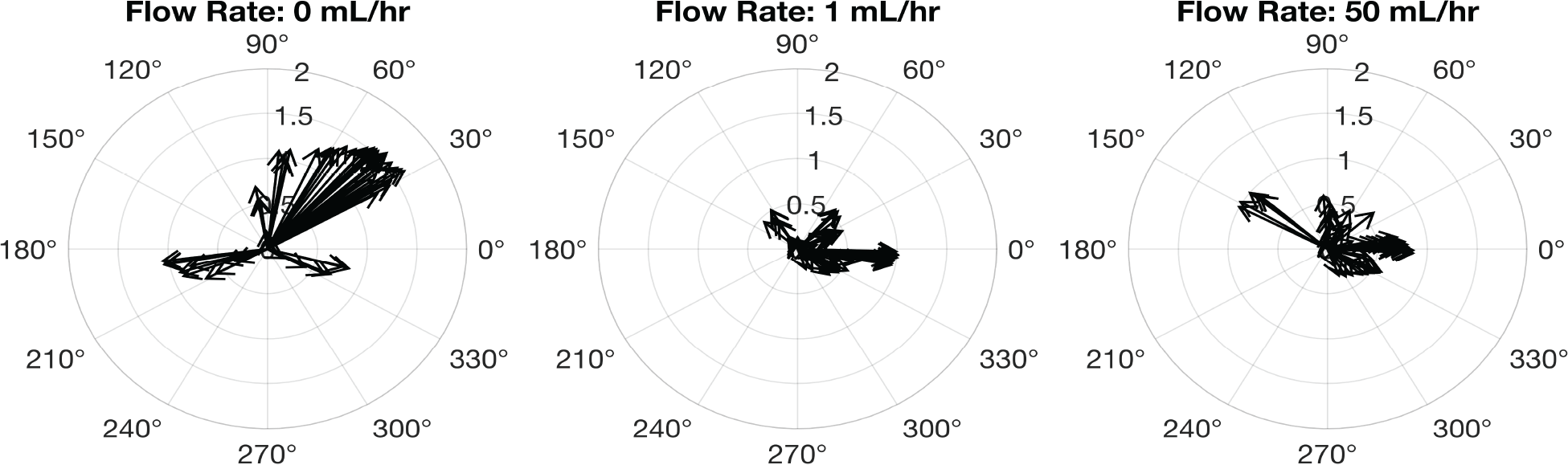
Sup Figure 2: *Hydra* in flow horizontally placed channels. For chambers positioned on a surface polyp displays a similar behavior; alignment with flow here is indicated as *θ* = 0 °.

**Figure 8:**
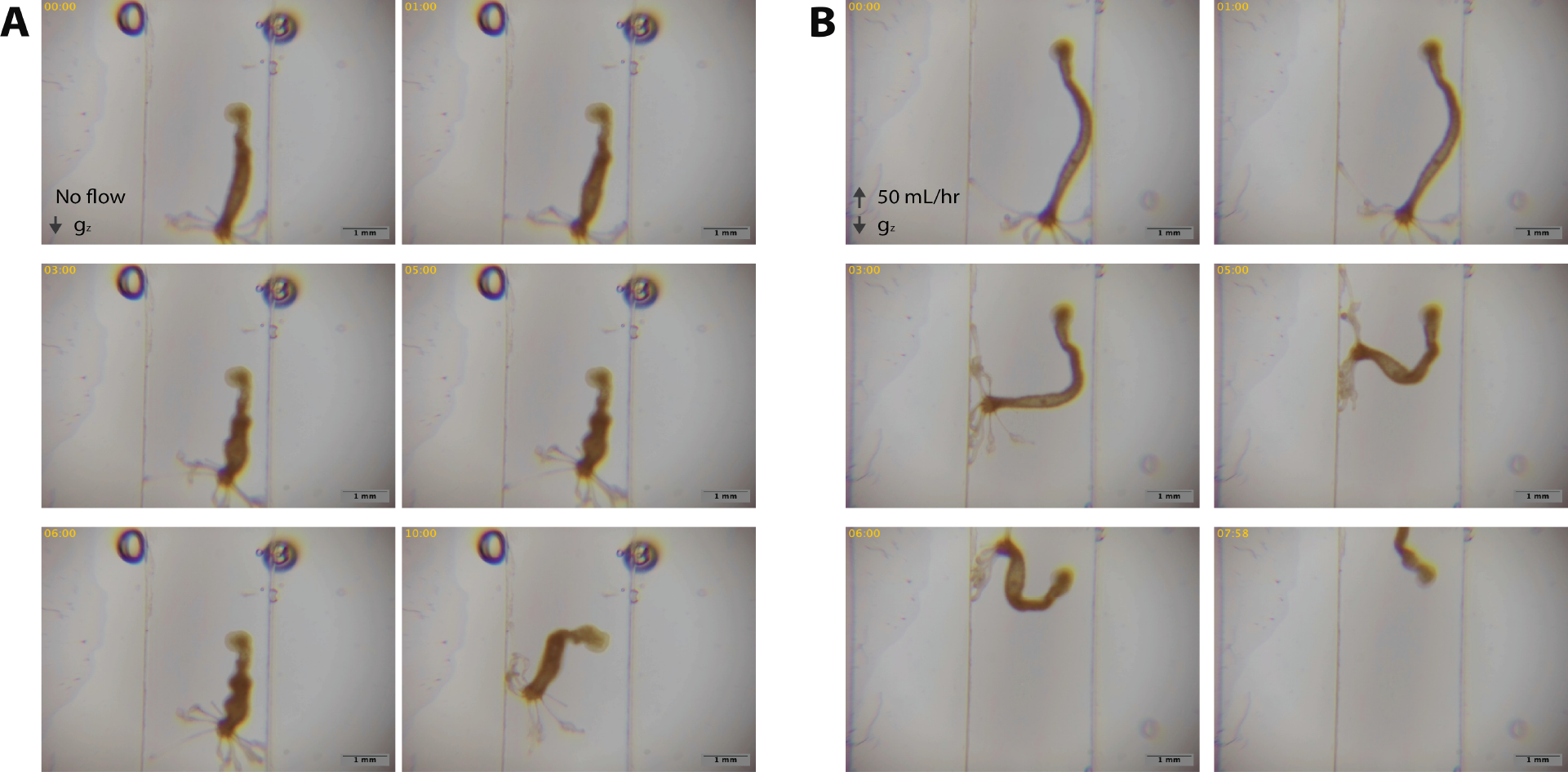
Sup Figure 3: *H. oxycnida* response to flow. (A) *H. oxycnida* remained surface attached without flow. (B) Polyp did not remain surface attached after the initiation of flow.

**Figure 9:**
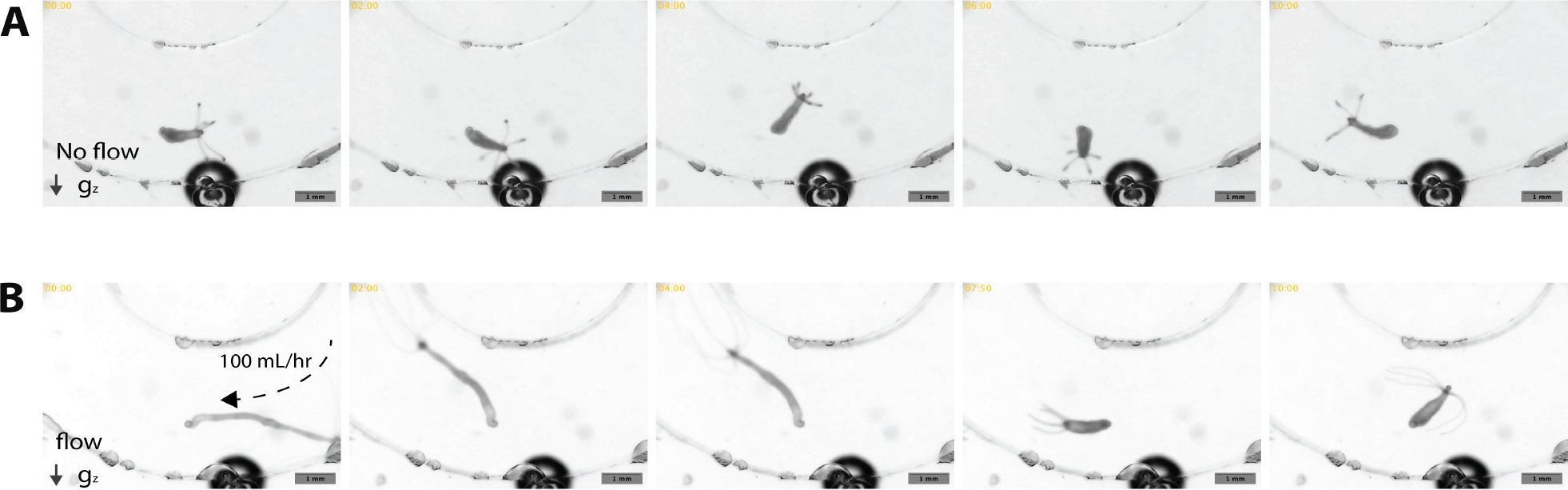
Sup Figure 4: Imaging *H. vulgaris* inside a curved channel. (A) without flow the polyp explored various directions. (B) At high flow rates (i.e., 100 mL/hr) the polyp aligns in the direction of the flow.

**Figure 10:**
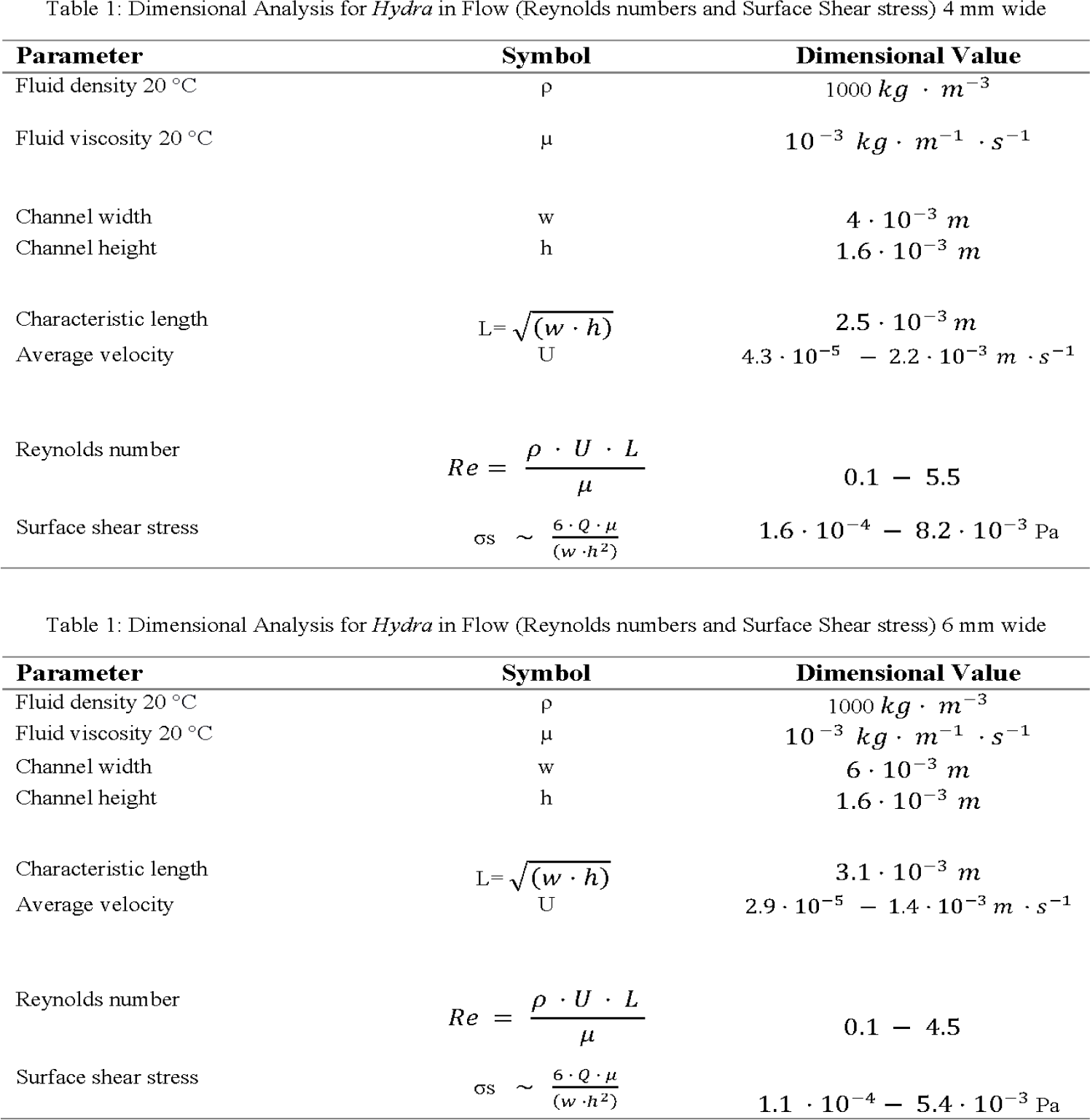
Table 1 Dimensional Analysis of the Chambers for Flow Conditions.

**Figure 11:**
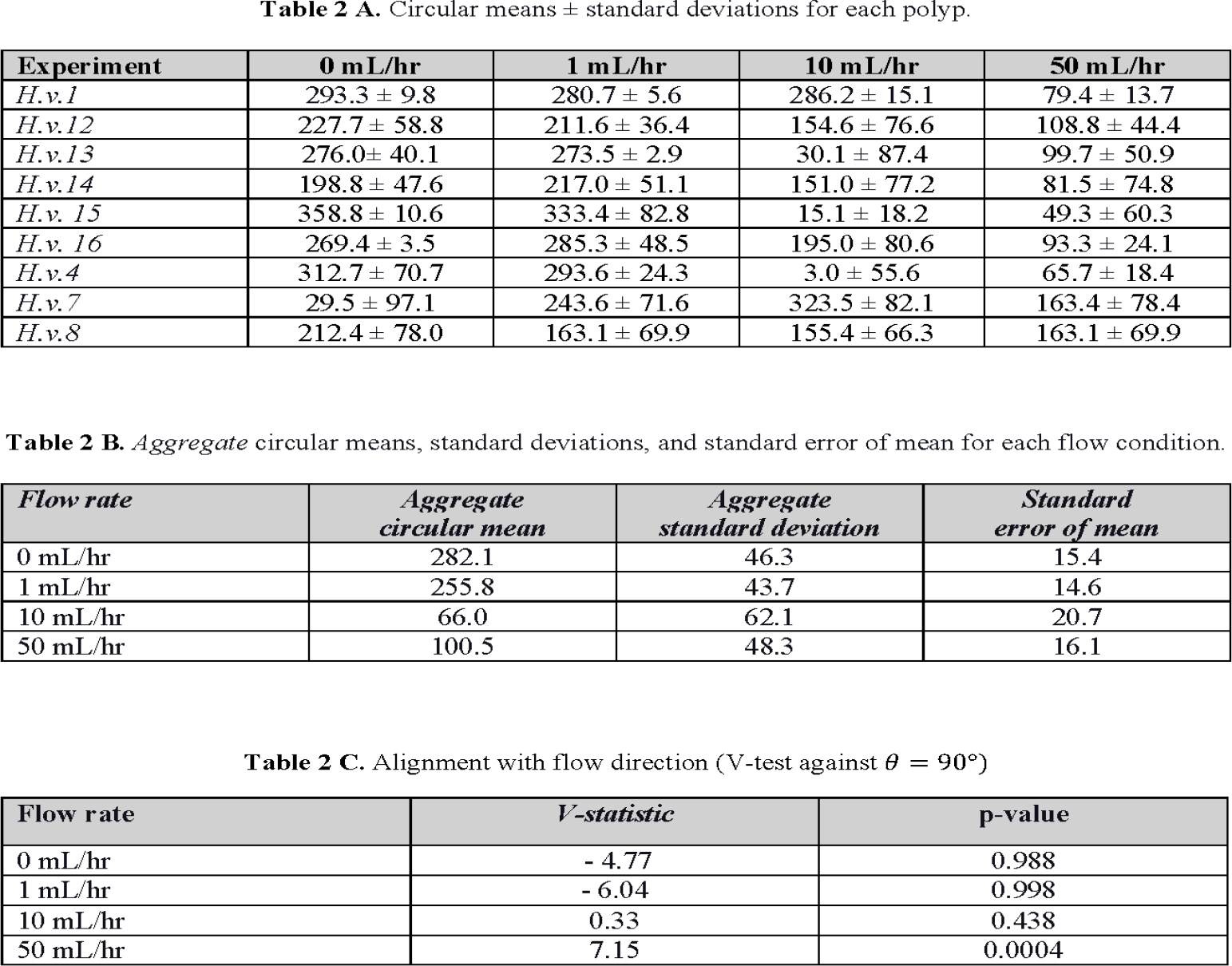
Table 2 Circular means and V-Test analysis.

Previous investigations, with in intact polyps and with tissue spheres, have established the connection between osmolarity and mechanosensing in *Hydra* (Kücken et al., 2008; Yamamoto et al., 2020; Ferenc et al., 2021) [13, 14, 15]. Specifically, results from Yamamoto et al. indicated that hypoosmotic conditions led to an increased hydrostatic pressure inside the *Hydra* resulting in whole body contractions (i.e., increase in mechanosensing). In contrast, in hyperosmotic solutions (relative to *Hydra*’s internal osmolarity) mechanosensing and contraction bursts were downregulated due to lower accumulation of water (Yamamoto et al., 2020)[14]. We focused on the combinatorial role of osmolarity and flow. To this aim, two organisms were exposed sequentially to 0 and 50 mL/hr flow of 0 Osm/L (for DI water) or a 50 mOsm/L (50 mM sucrose) solutions. To ensure that there were no effects due to the order of the solutions, both osmolarity sequences (high-to-low and low-to-high) were examined. As in previous experiments, the polyp remained surface attached in all conditions. Figure 4 provides these results indicating that changes in the osmolarity, and flow did not interfere with the surface attachment. In hyperosmotic solution (red vectors), organisms had reduced locomotion as compared to hypoosmotic solutions (blue vectors). This observation is in agreement with previous studies. Some alignment with flow was observed at high volumetric flow rates – particularly for the polyp that was first exposed to hyperosmotic solution. However, at this point the interplay between osmolarity and alignment with flow remains unclear.

Finally, we tested the effect of channel shape using a U-shaped curved channel (width 3.8 mm), to examine the response to flow in channels with a geometry different from the straight chambers. Adjusting the channel profile is valuable in introducing different velocity profiles. Here, the organism remained surface attached across various flow rates, and at high flow rates (here 100 mL/hr) it followed the direction of flow (Supplementary Figure 4).

## 4 Discussions

*Hydra* lives across diverse habitats, including still and moving water (Glauber et al., 2010)[31]. Given the organism’s presence in varied aquatic habitats, investigations of the role of environmental cues such as local mechanical stimuli (Rushforth et al., 1963; Badhiwala et al., 2021) [32, 17], osmotic pressure (Kücken et al., 2008; Wang et al., 2019; Yamamoto et al., 2020)[13, 33, 14], and light (Kim et al., 2024)[19] have been conducted. Additionally, as *H. vulgaris* has become a popular organism for biomechanics and neuroscience investigations (Carter et al., 2016; Wang et al., 2023; Yuste, 2024)[8, 34], studies have sought to elucidate the connections between sensorimotor biomechanics and the underlying neuronal activity [17]. We developed and verified resources to investigate the response of *Hydra* to flow. Moreover, by placing the chambers vertically, we sought to include the role of gravity. This choice was informed by an investigation that indicated a role for gravity (negative gravitotaxis), particularly for young buds (Ewer 1947) [4]. We experimented with the flow response of three different *Hydra* species. Overall, the variability in size and, to some degree, shape across the genus could provide excellent opportunities for biomechanics investigations. Yet, much of the established biomechanics work - including this report - has focused on *H. vulgaris*. This effort is the first report describing *H. hymanae* and *H. oxycnida* under flow conditions.

Using our microfluidic system, we examined two biomechanical responses of *H. vulgaris* to flow: surface attachment and body alignment under flow. Across the experiments, strong surface attachment was observed. Even the introduction of the fast-acting anesthetic 1 mM linalool (data not presented) with high flow (50 mL/hr) or adjusting the osmolarity of the solution did not lead to surface detachment (Goel et al., 2019)[35]. The role of secreted material for bioadhesion is known, and additionally, detachment may be initiated by mechanical forces via muscle contractions (Rodrigues et al., 2016)[6]. The presented strategy could provide an approach to investigate the adhesion/detachment across various surfaces (e.g., controlling for roughness or elastic modulus) or dissolved chemical factors that may interfere with adhesion. For *H. vulgaris*, we observed body column alignment with flow at high volumetric flow rates (50 mL/hr) for the majority of the polyps tested. Literature reports that the Young’s modulus of the upper body column of *Hydra* is approximately three times higher than that of the lower body column (Naik et al., 2020)[36]. The stiffness difference could explain the ease of stretching of the body column relative to the head. At this point, it remains unclear whether the flow alignment is an active or a passive process, or a mixture of both. A potential consequence of alignment with flow (independent of passive or active nature) could be related to the minimization of drag forces on the polyp. A change in the body shape (from cylindrical body to streamlined body) could dramatically change the drag coefficient and the corresponding drag force (Koehl 1982; Vogel 1996)[2, 37]. An interesting future direction is to investigate tradeoffs between flow alignment and prey capture. The relationship between transport (maximizing food capture) and minimizing mechanical forces in sea anemones including *Metridium senile* (Koehl 1977)[3] and *Alcyonium siderium* (Patterson 1991)[38] have been investigated. Highlighting the size and shape differences, *Hydra* may experience a similar relationship between flow regime and prey capture. Indeed, while *Hydra* in still water may not be exposed to prolonged flow, during short bursts of rapid flow, surface attachment could be more critical than feeding. We also note that we did not observe a change in contraction bursts across different conditions. However, longer recordings for a specific condition may be needed to explore a potential relationship between the number of contractions and flow.

Future investigations could study the molecular and neuronal basis of flow response via the existing transgenic lines expressing calcium reporters (Dupre et al., 2017) [11] and pharmacological agents (linalool or reduced glutathione). Specifically, recent investigations suggest a role for mechanosensitive ion channels in the behavior of *Hydra* (Skokan et al., 2024; Alzugaray et al., 2025)[39, 40]. Combining the pharmacological agents with flow under well-controlled microfluidic chambers could provide exciting future opportunities.

## Acknowledgments

We are grateful to Professor Bruno Pernet for his support, valuable discussions, and feedback. Authors are thankful to CSULB Marine Lab, specifically Yvette Ralph. Authors are thankful to the members of the Ahrar-lab specifically: B.A. Lopez, H. Nandasena, H.B. Obenshain, and K. Yoshida for their support of the project.

Co-authors indicated by ⋆ equally contributed to the project.

The authors gratefully acknowledge financial support for this research by a grant from the STEM-NET consortium at the CSU Office of the Chancellor. This work was also supported in part by: (a) A CSUPERB new investigator award to SA; (b) the CSULB BUILD Program in the Office of Undergraduate Research Services; (c) US Department of Education, Office of Postsecondary Education (OPE), Asian American and Native American Pacific Islander-Serving Institutions (AANAPISI) Program under Award Numbers P031L220020-AANAPISI Part A.

## Affiliations

- 1: Department of Biomedical Engineering. California State University Long Beach, CA, US.
- 2: Department of Biological Chemistry, University of California Irvine, Irvine, CA 92697, USA

## OSF Repository

As part of the project, resources, including the design files, and code for analysis, have been shared.

OSF Name: *Hydra* in Flow.

Link: will be added.

## Institutional Review Board Statement

Not applicable.

## List of Figures

- Figure 1: Preliminary experiments suggested alignment between *Hydra vulgaris* and flow at high volumetric flow rates.
- Figure 2: Three polyps in flow.
- Figure 3: Overall assessment of alignment with flow as depicted by circular means for each polyp.
- Figure 4: *Hydra hymanae* response to flow.
- Figure 5: Flow and osmotic pressure.

## List of Supplementary Figures

- Sup Figure 1: *H. vulgaris* polyp in flow (0 - 100 mL/hr).
- Sup Figure 2: *H. vulgaris* in flow inside a horizontal channel (no gravity).
- Sup Figure 2: *H. oxycnida* in flow.
- Sup Figure 3: *H. vulgaris* inside a curved channel

## List of Supplementary Movies

- Movie 1: *H. vulgaris* in flow.
- Movie 2: *Hydra hymanae* in flow.

## List of Tables

- Table 1: Dimensional analysis of chamber properties.
- Table 2: Summary of responses to flow.

## References

[1] D. Martínez, A. Iñiguez, KM Percell, JB Willner, J Signorovitch, and RD Campbell. Phylogeny and biogeography of hydra (cnidaria: Hydridae) using mitochondrial and nuclear dna sequences. Molecular Phylogenetics and Evolution, 57(1): 403–410, 2010.

[2] MAR Koehl. The interaction of moving water and sessile organisms. Scientific American, 247(6): 124–135, 1982.

[3] MAR Koehl. Effects of sea anemones on the flow forces they encounter. Journal of Experimental Biology, 69(1): 87–105, 1977.

[4] RF Ewer. Tne behaviour of hydra in response to gravity. In Proceedings of the zoological Society of London, volume 117, pages 207–218. Wiley Online Library, 1947.

[5] Matthias C Vogg, Brigitte Galliot, and Charisios D Tsiairis. Model systems for regeneration: Hydra. Development, 146(21), 2019.

[6] Marcelo Rodrigues, Philippe Leclère, Patrick Flammang, Michael W Hess, Willi Salvenmoser, Bert Hobmayer, and Peter Ladurner. The cellular basis of bioadhesion of the freshwater polyp hydra. BMC Zoology, 1(1): 3, 2016.

[7] Richard D Campbell. Structure of the mouth of hydra spp. a breach in the epithelium that disappears when it closes. Cell and tissue research, 249(1): 189–197, 1987.

[8] Jason A Carter, Callen Hyland, Robert E Steele, and Eva-Maria S Collins. Dynamics of mouth opening in hydra. Biophysical journal, 110(5): 1191–1201, 2016.

[9] John R Szymanski and Rafael Yuste. Mapping the whole-body muscle activity of hydra vulgaris. Current Biology, 29(11): 1807–1817, 2019.

[10] Athina Keramidioti, Sandra Schneid, Christina Busse, Christoph Cramer von Laue, Bianca Bertulat, Willi Salvenmoser, Martin Hess, Olga Alexandrova, Kristine M Glauber, Robert E Steele, et al. A new look at the architecture and dynamics of the hydra nerve net. Elife, 12:RP87330, 2024.

[11] Christophe Dupre and Rafael Yuste. Non-overlapping neural networks in hydra vulgaris. Current Biology, 27(8): 1085–1097, 2017.

[12] Hengji Wang, Joshua Swore, Shashank Sharma, John R Szymanski, Rafael Yuste, Thomas L Daniel, Michael Regnier, Martha M Bosma, and Adrienne L Fairhall. A complete biomechanical model of hydra contractile behaviors, from neural drive to muscle to movement. Proceedings of the National Academy of Sciences, 120(11):e2210439120, 2023.

[13] Michael Kücken, Jordi Soriano, Pramod A Pullarkat, Albrecht Ott, and Ernesto M Nicola. An osmoregulatory basis for shape oscillations in regenerating hydra. Biophysical journal, 95(2): 978–985, 2008.

[14] Wataru Yamamoto and Rafael Yuste. Whole-body imaging of neural and muscle activity during behavior in hydra vulgaris: effect of osmolarity on contraction bursts. eneuro, 7(4), 2020.

[15] Jaroslav Ferenc, Panagiotis Papasaikas, Jacqueline Ferralli, Yukio Nakamura, Sebastien Smallwood, and Charisios D Tsiairis. Mechanical oscillations orchestrate axial patterning through wnt activation in hydra. Science advances, 7(50):eabj6897, 2021.

[16] Janna C Nawroth, Christoph Giez, Alexander Klimovich, Eva Kanso, and Thomas CG Bosch. Spontaneous body wall contractions stabilize the fluid microenvironment that shapes host–microbe associations. Elife, 12:e83637, 2023.

[17] Krishna N Badhiwala, Abby S Primack, Celina E Juliano, and Jacob T Robinson. Multiple neuronal networks coordinate hydra mechanosensory behavior. elife, 10:e64108, 2021.

[18] Wataru Yamamoto and Rafael Yuste. Peptide-driven control of somersaulting in hydra vulgaris. Current Biology, 33(10): 1893–1905, 2023.

[19] Soonyoung Kim, Krishna N Badhiwala, Guillaume Duret, and Jacob T Robinson. Phototaxis is a satiety-dependent behavioral sequence in hydra vulgaris. Journal of Experimental Biology, 227(18):jeb247503, 2024.

[20] Shuting Han, Ekaterina Taralova, Christophe Dupre, and Rafael Yuste. Comprehensive machine learning analysis of hydra behavior reveals a stable basal behavioral repertoire. Elife, 7:e32605, 2018.

[21] Krishna N Badhiwala, Daniel L Gonzales, Daniel G Vercosa, Benjamin W Avants, and Jacob T Robinson. Microfluidics for electrophysiology, imaging, and behavioral analysis of hydra. Lab on a Chip, 18(17): 2523–2539, 2018.

[22] Constantine N Tzouanas, Soonyoung Kim, Krishna N Badhiwala, Benjamin W Avants, and Jacob T Robinson. Hydra vulgaris shows stable responses to thermal stimulation despite large changes in the number of neurons. Iscience, 24(6), 2021.

[23] Thomas Perros, Anaïs Biquet-Bisquert, Zacchari Ben Meriem, Morgan Delarue, Pierre Joseph, Philippe Marcq, and Olivier Cochet-Escartin. Mechanical characterization of regenerating hydra tissue spheres. Biophysical Journal, 123(13): 1792–1803, 2024.

[24] Per Niklas Hedde, Brian T Le, Erika L Gomez, Leora Duong, Robert E Steele, and Siavash Ahrar. Spim-flow: an integrated light sheet and microfluidics platform for hydrodynamic studies of hydra. Biology, 12(1): 116, 2023.

[25] Neha Khetan, Shagun Maheshwari, and CAA Athale. Quantitative detachment mechanics of hydra from substrates. In Proc. Indian Natn. Sci. Acad, volume 85, pages 291–299, 2019.

[26] Haley B Obenshain, Isaias Zarate, Olivia Hedman-Manzano, Jared Goderich, Sungho Lee, Bryant A Lopez, Emma Varela, Ga-Young Kelly Suh, Douglas A Pace, and Siavash Ahrar. Aquavert–imaging and microfluidics for vertical swimming of microorganisms. bioRxiv, pages 2024–09, 2024.

[27] Charles Elmer Hadley, Helen Forrest, et al. Taxonomic studies on the hydras of north america. 6, description of hydra hymanae, new species. american museum novitates; no. 1423. 1949.

[28] Johannes Schindelin, Ignacio Arganda-Carreras, Erwin Frise, Verena Kaynig, Mark Longair, Tobias Pietzsch, Stephan Preibisch, Curtis Rueden, Stephan Saalfeld, Benjamin Schmid, et al. Fiji: an open-source platform for biological-image analysis. Nature methods, 9(7): 676–682, 2012.

[29] David Long. True circular mean. MATLAB Central File Exchange, 2023,. (https://www.mathworks.com/matlabcentral/fileexchange/132118-true-circular-mean); Retrieved August 19, 2025.

[30] Philipp Beren. Circular statistics toolbox (directional statistics). MATLAB Central File Exchange, 2025. (https://www.mathworks.com/matlabcentral/fileexchange/10676-circular-statistics-toolbox-directional-statistics); Retrieved August 19, 2025.

[31] Kristine M Glauber, Catherine E Dana, and Robert E Steele. Hydra. Current Biology, 20(22):R964–R965, 2010.

[32] Norman B Rushforth, Allison L Burnett, and Richard Maynard. Behavior in hydra: contraction responses of hydra pirardi to mechanical and light stimuli. Science, 139(3556): 760–761, 1963.

[33] Rui Wang, Tapan Goel, Kate Khazoyan, Ziad Sabry, Heng J Quan, Patrick H Diamond, and Eva-Maria S Collins. Mouth function determines the shape oscillation pattern in regenerating hydra tissue spheres. Biophysical Journal, 117(6): 1145–1155, 2019.

[34] Rafael Yuste. Breaking the neural code of a cnidarian: Learning principles of neuroscience from the “vulgar” hydra. Current Opinion in Neurobiology, 86:102869, 2024.

[35] Tapan Goel, Rui Wang, Sara Martin, Elizabeth Lanphear, and Eva-Maria S Collins. Linalool acts as a fast and reversible anesthetic in hydra. PloS one, 14(10):e0224221, 2019.

[36] Suyash Naik, Manu Unni, Devanshu Sinha, Shatruhan Singh Rajput, Puli Chandramouli Reddy, Elena Kartvelishvily, Inna Solomonov, Irit Sagi, Apratim Chatterji, Shivprasad Patil, et al. Differential tissue stiffness of body column facilitates locomotion of hydra on solid substrates. Journal of Experimental Biology, 223(20):jeb232702, 2020.

[37] Steven Vogel. Life in moving fluids: the physical biology of flow. Princeton university press, 1996.

[38] Mark R Patterson. The effects of flow on polyp-level prey capture in an octocoral, alcyonium siderium. The Biological Bulletin, 180(1): 93–102, 1991.

[39] Taylor D Skokan, Bert Hobmayer, Kara L McKinley, and Ronald D Vale. Mechanical stretch regulates macropinocytosis in hydra vulgaris. Molecular Biology of the Cell, 35(3):br9, 2024.

[40] María Eugenia Alzugaray, María Victoria Gavazzi, Lucía Griffo, and Jorge Rafael Ronderos. Piezo proteins, mechano reception and behaviour in hydra. Scientific Reports, 15(1): 6440, 2025.

